# miRNA mediated downregulation of non-muscle Cyclase associated protein 1 is required for myogenic differentiation

**DOI:** 10.1101/2021.10.26.465871

**Authors:** Anurag Kumar Singh, Amrita Rai, Anja Weber, Guido Posern

## Abstract

Myoblast fusion is crucial for the formation, growth and regeneration of healthy skeletal muscle, but the molecular mechanisms that govern fusion and myofiber formation remain poorly understood. Here we report that Cyclase-associated protein 1 (Cap1), a regulator of actin dynamics, plays a critical role in cytoskeletal remodeling during myoblast fusion and formation of myotubes. Cap1 mRNA and protein are expressed in murine C2C12 and human LHCN-M2 myoblasts, but its abundance decreases during myogenic differentiation. Perturbing the temporally controlled expression of Cap1 by overexpression or Crispr-Cas9 mediated knockout impaired actin rearrangement, myoblast alignment, expression of profusion molecules, differentiation into multinucleated myotubes and myosin heavy chain expression. Endogenous Cap1 expression is posttranscriptionally downregulated during differentiation by canonical myomiRs miR-1, miR-133 and miR-206, which have conserved binding sites in the 3’ UTR of the Cap1 mRNA. Deletion of the endogenous 3’ UTR in C2C12 cells phenocopies overexpression of Cap1 by inhibiting myotube formation. Our findings implicate Cap1 and its myomiR-mediated downregulation in the myoblast fusion process and the generation of skeletal muscle.

## Introduction

Skeletal muscle is the largest tissue in the body, accounting for ~40% of human body mass. A fundamental step in the late differentiation process of muscle cells is the fusion of mononucleated myoblasts to multinucleated myofibers (*1–3*). Similarly, in response to injury, myogenic progenitor cells within the adult musculature are activated and fuse to regenerate myofibers. Many studies have provided insights regarding the mechanisms and molecular components that mediate skeletal muscle myoblast fusion. These include identification of the proteins mediating cell-cell adhesion and recognition of pathways that relay fusion signals from the cell membrane to the cytoskeleton (*1, 2*). Despite these insightful studies, a complete understanding of the mechanisms governing the fusion process is lacking.

Dramatic reorganization of the cytoskeleton occurs as myoblasts maneuver through the morphological changes associated with cell-cell fusion to form multinucleated myotubes. These morphological changes include myoblast migration, elongation to a bipolar shape, membrane alignment and fusion (*1, 3*). In the live embryos dynamic F-actin foci found at the point of cell-cell contact are formed and dissolves, which coincided with myoblast fusion event (*4, 5*). In flies, a dense actin focus in fusion-competent myoblasts invades into founder cells through a thin sheath of F-actin, which requires the action of the nucleation-promoting factors (NPFs) Scar and WASP on the Arp2/3 complex (*6, 7*). Similar to flies, extensive cytoskeletal reorganization occurs before and after fusion in cultured mammalian myoblasts. Visualization of the F-actin cytoskeleton revealed dynamic changes in fusing mammalian myoblasts *in vitro* and a dense F-actin wall paralleling the long axis of aligned myoblasts (*8–10*). As fusion proceeds, gaps appear in this actin wall at sites of vesicle accumulation, and the fusion pores form. In the absence of proper remodeling, F-actin structures continuously accumulate at the site of cell-cell contact and are correlated with a decrease in myoblast fusion (*9*).

Cyclase-associated proteins (CAP) are evolutionarily conserved proteins with largely unknown physiological functions. Early studies suggested a rather passive role for CAP in actin cytoskeleton regulation, which was believed to act via sequestering globular actin monomers (*11*). This view has changed considerably in the last decade because CAP has been implicated in almost all steps relevant for actin dynamics. Specifically, these studies unraveled (i) a co-operation of CAP with key actin regulators such as ADF/Cofilin and Twinfilin in F-actin disassembly, (ii) a nucleotide exchange activity on G-actin that is required for F-actin assembly, and (iii) an inhibitory function towards the F-actin assembly factor inverted formin 2 (INF2) (*12–15*). Unlike yeast, mammals possess two Cap family members, Cap1 and Cap2, with different expression patterns. CAP2 is abundant in the heart, striated muscle, and brain and is required for skeletal muscle development, heart physiology, and synaptic function (*16–19*). Instead, CAP1 expression is less restricted, but its physiological functions remain largely unclear, due to the lack of an appropriate mouse model (*20*), while a recent brain-specific Cap1 KO mouse study shows its role in the control of neuronal actin dynamics and growth cone morphology together with Cofilin1 (*21*). Although embryonic expression in murine muscle has been observed for Cap1, it decreases postpartum to undetectable levels in the adult and has not yet been studied in relation to skeletal muscle development or differentiation (*22, 23*).

In this study, we report a downregulation in Cap1 expression during the differentiation of murine C2C12 and human LHCN-M2 cells from myoblast to myotubes. Loss- and gain-of-function experiments show that expression of Cap1 in myoblasts, as well as its differentiation-induced downregulation, is required for myotube formation. Further, we show that the downregulation of the Cap1 expression is controlled via miRNA-mediated degradation of the Cap1 mRNA, which results in the loss of CAP1 protein. We conclude that the timely loss of CAP1 is an important event during myoblast fusion into myotubes.

## Results

### Cap1 mRNA and protein levels are downregulated during myogenic differentiation

Both murine, as well as human skeletal muscle cell lines C2C12 and LHCN-M2, are able to recapitulate terminal differentiation of myoblasts to fused, multinuclear myotubes. To elucidate this process, cells were seeded at high density, and differentiation was induced by changing to a differentiation medium. The first multinucleated myotubes were observable at day 4 (Fig. 1A and E, d4) with more mature myotubes at day 6 (Fig. 1A and E, d6). Subsequently, the differentiation process was verified by the appearance and subsequent upregulation of myosin heavy chain polypeptides MYH1, MYH2, MYH4, and MYH6 over the course of differentiation (Fig. 1C and G).

**Fig. 1:**
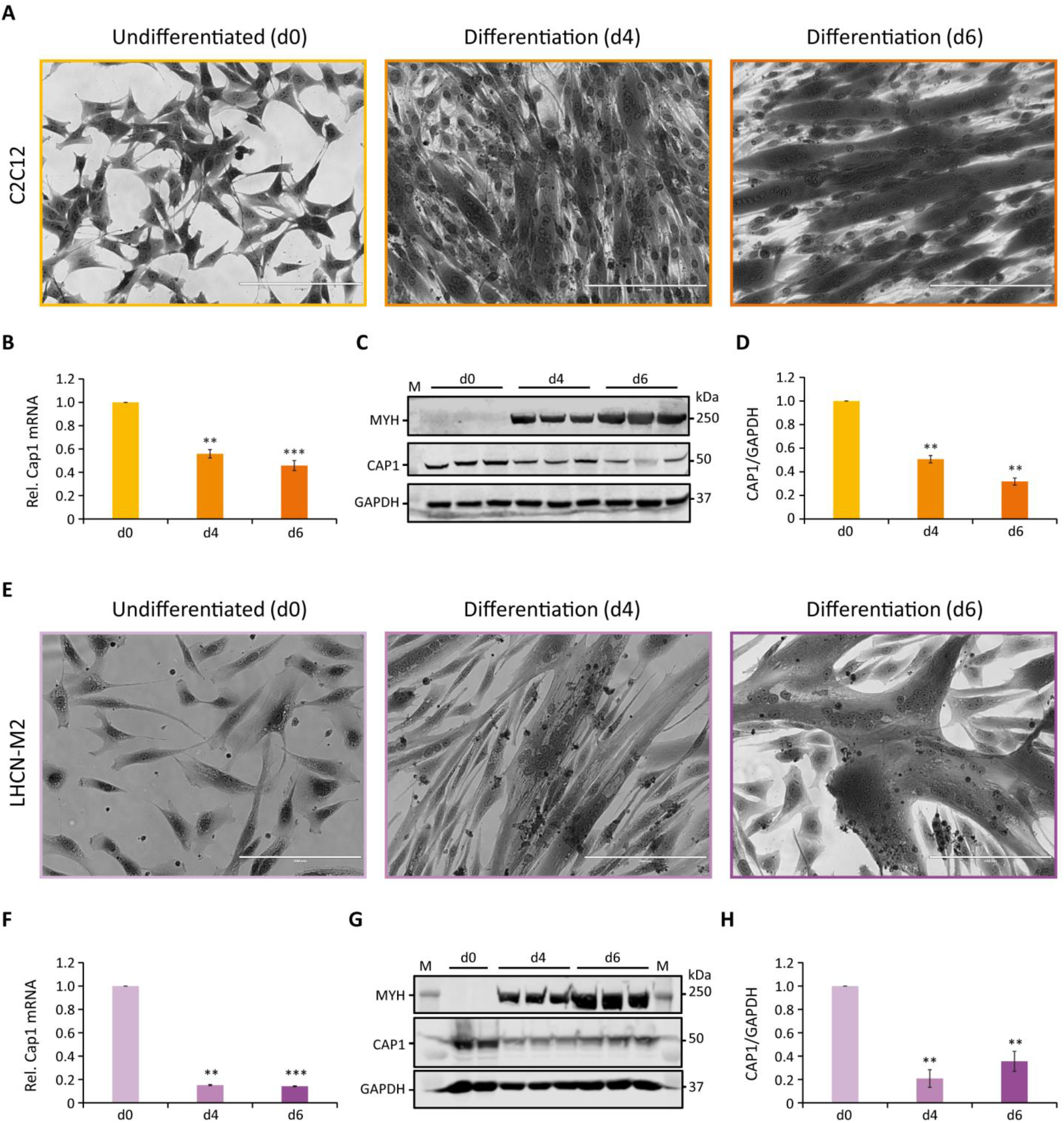
Cap1 mRNA and protein levels are downregulated during myogenic differentiation. (**A** and **E**) Bright-field images (20x) of murine C2C12 (**A**) and human LHCN-M2 (**E**) cells upon differentiation for 4 and 6 days (d4 and d6), in comparison to undifferentiated control cells (d0). Cells were stained with crystal violet. (**B** and **F**) Relative Cap1 mRNA in differentiating C2C12 (**B**) and LHCN-M2 (**F**) cells, quantified by qRT-PCR and normalized to a set of housekeeping mRNAs. (**C** and **G**) Immunoblots of lysates from differentiating C2C12 (**C**) and LHCN-M2 (**G**) cells, using antibodies against myosin heavy chain polypeptides 1, 2, 4 and 6 (MYH), CAP1 and GAPDH as a control. (**D** and **H**) Quantification of CAP1 immunoblots at myogenic differentiation for 4 and 6 days, normalized to undifferentiated control cells. *Error bars*, SEM (n = 3); **p< 0.01, ***p< 0.001 (Student’s t-test). Scale bar, 200 μm.

CAP proteins have been implicated in the regulation of actin treadmilling and the differentiation of muscle cells. Because of the importance of actin remodeling during myoblast fusion, we analyzed whether or not CAP1 plays any role in myogenic differentiation. Muscle expression of Cap1 at the embryonic stage had been documented, but it decreases postpartum and is undetectable in adult mice (*22*). We thus assessed Cap1 expression during C2C12 and LHCN-M2 differentiation both at mRNA as well as protein level. Cap1 mRNA and protein were clearly expressed in undifferentiated cells but decreased upon differentiation (Fig. 1 B, C, D, F, G, H). In contrast, Cap2 mRNA was hardly detectable in undifferentiated myoblasts but was upregulated in differentiating cells, consistent with its previously described role in postnatal skeletal muscle development (Fig. S1)(*19*). To further validate the inverse regulation of Cap1 and Cap2 during myogenic differentiation and regeneration, we re-analyzed two existing microarray datasets from the GEO database (*24*). The first one investigated the genome-wide mRNA expression of C2C12 cells differentiating from myoblasts to myotubes (GSE4694) (*25*). The second one analyzed the gene expression in the mouse tibialis anterior (TA) muscle of 12 wk old C57BL/6J males after injury by glycerol injection for up to 21 days (GSE45577) (*26*). The data analysis revealed a significant upregulation of Cap1 immediately post TA injury, with subsequent downregulation towards 21 days as the injury heals and the muscle is regenerated *in vivo* (Fig. S2). Consistent with this and our data, Cap1 is also expressed in the dataset of C2C12 myoblasts, but downregulated in myotube formation *in vitro* (Fig. S3) (*25*). In contrast, both studies find increased levels of Cap2 upon differentiation *in vitro* or regeneration *in vivo*, in line with our observation of inversely regulated Cap1 and Cap2 mRNA in differentiating C2C12 or LHCN-M2 cells. These data demonstrate Cap1 expression in myoblasts and suggest a role for timely downregulation upon differentiation.

### Cap1 deletion results in increased cell size and F-actin organization

To investigate whether the observed decrease in Cap1 expression plays a role in myogenic differentiation, we perturbed the expression of Cap1 and assessed if that affects the myoblast fusion and myogenic differentiation. First, we analyzed whether or not the knockout (KO) of Cap1 has any impact on F-actin. Knockout of Cap1 in C2C12 cells by two sgRNA using CRISPR-Cas9 and subsequent selection generated a population of cells with reduced CAP1 protein abundance (Fig. 2A). This resulted in profoundly increased cell size, a spread-out morphology and enhanced F-actin filament formation, resulting in stable stress fiber (Fig. 2B, C; Fig. S4). This result is consistent with the role of CAP1 as an F-actin depolymerizing protein (*14, 15*). In contrast, cells overexpressing Cap1 (Flag-tagged dsRed-CAP1) did not look much different from the undifferentiated Cas9 control cells, and quantification of the cell area covered showed no significant changes (Fig. 2B; Fig. S4).

**Fig. 2:**
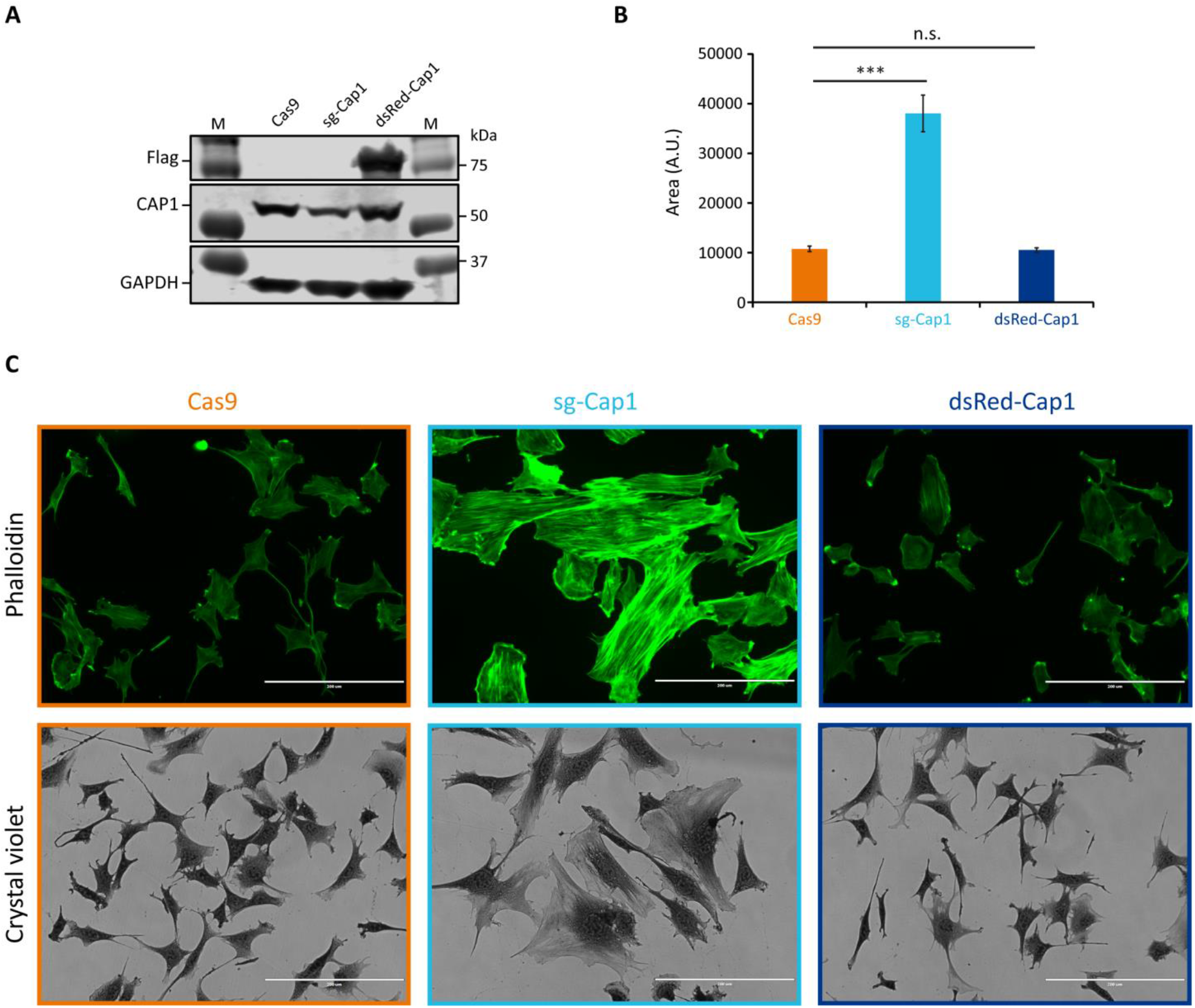
Knockout of Cap1 results in increased cell size and accumulated F-actin fibers in C2C12 cells. (**A**) Validation of partial knockout by CRISPR-Cas9 (sg-Cap1) and overexpression (dsRed-Cap1) in C2C12 pools by immunoblot, compared to control cells infected with Cas9 only (Cas9). The sg-Cap1 cells show reduced expression of endogenous CAP1 while ectopically expressed dsRed-CAP1 results in an additional band corresponding to tagged CAP1 proteins. (**B**) Quantification of the cell area covered, based on crystal violet staining. (**C**) Micrographs. The upper panel shows fluorescence images of phalloidin-stained cells. The lower panel shows bright-field images (20x) of crystal violet stained cells. *Error bars*, SEM (n= 50 cells); ***p< 0.001 (Student’s t-test). Scale bar, 200 μm.

### Deregulated Cap1 expression inversely correlates with myogenic differentiation

F-actin reorganization is critical for myoblast fusion and myogenic differentiation. To test whether Cap1 downregulation marks an important event, we studied the differentiation process of the C2C12 cells with perturbed expression of Cap1. Under both circumstances (Cap1 overexpression and knock-out), the differentiation was severely affected (Fig. 3; Fig. S5). On day 2, the control cells were spindle-shaped and aligned to each other (Fig. S5, Cas9). The Cap1 KO cells seemed bigger and more elongated, while the Cap1 overexpressing cells lacked this phenotype at day 2 and maintained a fibroblast-like phenotype, similar to undifferentiated cells (Fig. S5, d2). Both the overexpression and the knockout of the Cap1 changed the way cells align to each other at the initial phase of differentiation and affect the fusion process. The result suggests an important role for the timely downregulation of the CAP1 protein during the differentiation process. In accordance, cell movement and change in cell shape followed by cell-cell adhesion are known to be important steps during myoblast fusion and myogenic differentiation.

**Fig. 3:**
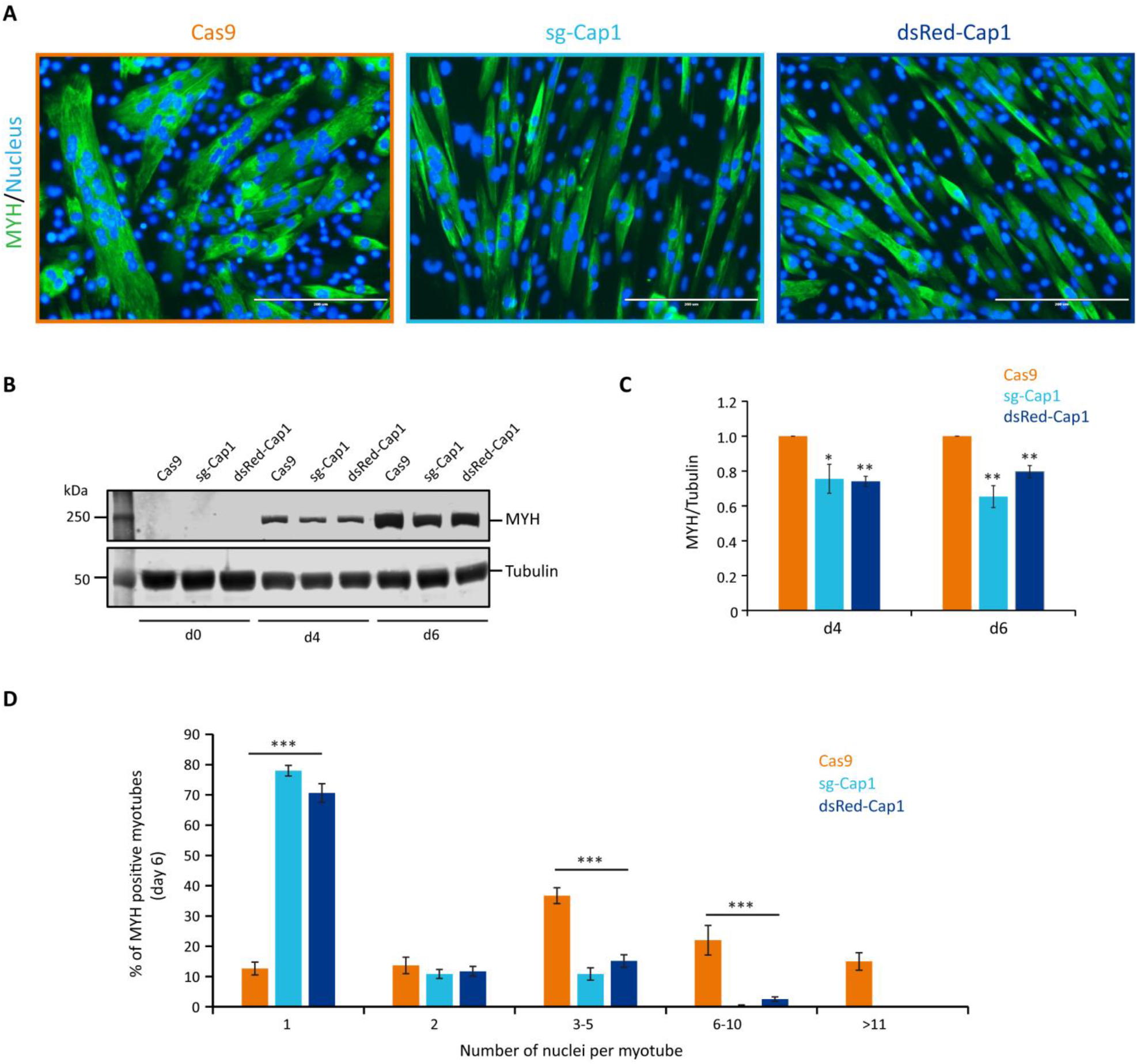
Timely downregulation of Cap1 is important for proper myogenic differentiation. (**A**) Cas9 control cells, knockout (sg-Cap1) and overexpressing (dsRed-Cap1) cells at day 6 of their differentiation. Pools of C2C12 cells were stained for MYH (green) and nucleus (DAPI, blue). (**B**) Western blot for the myogenic marker MYH at days 4 and 6 of the differentiation. (**C**) Quantification of MYH immunoblots at myogenic differentiation for 4 and 6 days, normalized to Cas9 control cells. (**D**) Quantification of the percentage of myotubes containing the indicated number of nuclei per myotube in control, knockout and overexpression myotubes after 6 days of differentiation. *Error bars*, SEM (minimum 200 MYH positive cells were counted); * p< 0.05, ** p< 0.01, ***p< 0.001 (Student’s t-test). Scale bar, 200 μm.

Further, we analyzed the expression of the differentiation marker myosin heavy chain (MYH) by immunofluorescence microscopy (Fig. 3A) and western blot analysis (Fig. 3B, C). Compared to the Cas9 control cells, the MYH positive myotubes appeared thinner and with less number of nuclei per myotubes upon Cap1 knockout and overexpression (Fig. 3A, D). Western blot analysis confirmed that both overexpression and Cap1 knockout resulted in decreased levels of MYH over the course of differentiation (Fig. 3B and C).

Next, we analyzed the number of nuclei present in the MYH-positive myotubes at day 6 of differentiation. Cas9 control myotubes showed significantly higher numbers of nuclei compared to both the knockout as well as Cap1 overexpressing cells (Fig. 3D). In contrast, the majority of MYH-positive “myotubes” with altered Cap1 expression contained only one nucleus. This points towards a defect in the fusion process during the differentiation. Since the expression of Cap1 was decreasing upon differentiation (Fig. 1), and perturbation of Cap1 expression impaired differentiation, we conclude that a timely decrease of Cap1 is paramount to myoblast fusion and maturation.

### Cap1 knockout, as well as overexpression, resulted in aberrant pre-fusion F-actin organization

During differentiation of myoblast to myotubes, the actin filaments undergo dramatic reorganization. In the mammalian system, it has been shown that myoblasts develop a non-uniform thickened actin wall as a prerequisite for the fusion event (*8*). Thus we looked for changes in the F-actin organization after one day in differentiation medium and whether it was affected by altering the expression of Cap1 in C2C12 cells. Compared to Cas9 control cells which show F-actin bundles at the lateral sides of the aligned spindle-shaped cells (Fig. 4A, arrows), Cap1 knockout cells (sg-Cap1) showed the frequent appearance of F-actin foci and thick disorganized accumulation of F-actin (Fig. 4A, arrowheads). In contrast, Cap1 overexpressing cells (dsRed-Cap1) lacked the elongated cell phenotype and the F-actin organization appeared similar to the undifferentiated cells, consistent with a delayed response to the differentiation medium.

**Fig. 4:**
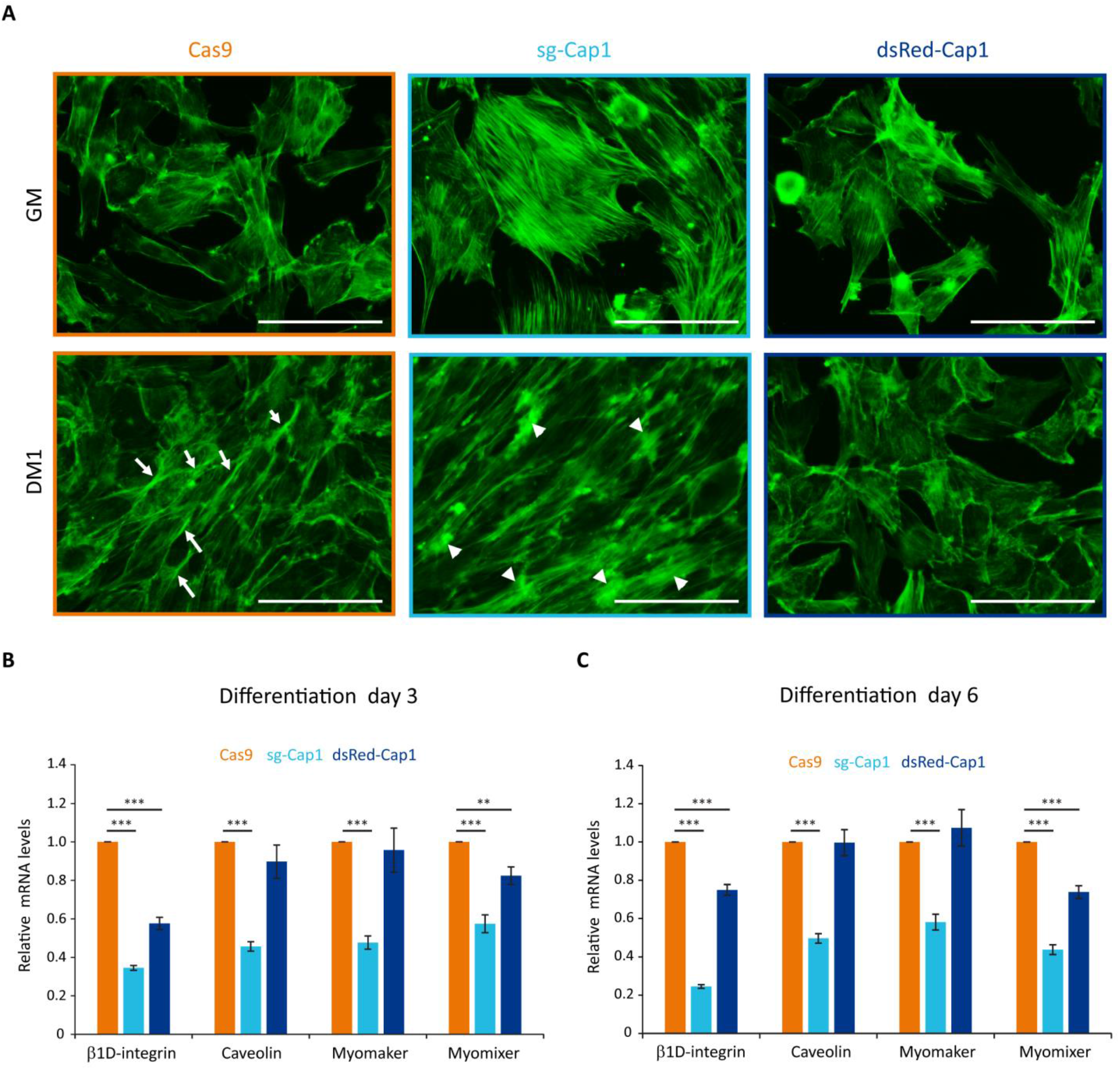
Disturbance of the early cortical actin rearrangement and expression of pro-fusion molecules upon changes in Cap1 expression. (**A**) Cas9 control, sg-Cap1 and dsRed-Cap1 cells were cultured in growth medium (GM) or differentiation medium (DM1; day 1 of differentiation) and fixed and stained to detect F-actin using phalloidin. Aligned Cas9 control cells (lower panel) show longitudinal actin fiber accumulation at sites of contact (arrows), whereas Cap1 knockout cells (sg-Cap1, middle-lower panel) exhibit mislocalized, thickened actin patches (arrowheads). Alignment and cortical actin fibers are absent in dsRed-Cap1 cells (last panel). Scale bar, 100 μm. (**B** and **C**) The mRNAs for ß1D-integrin, caveolin, myomaker and myomixer were quantified by qRT-PCR and normalized to a set of housekeeping mRNAs at day 3 (**B**) and day 6 (**C**) of differentiation. *Error bars*, SEM (n = 3); **p< 0.01, *** p< 0.001 (Student’s t-test).

### Perturbation in Cap1 expression resulted in the downregulation of pro-fusion molecules

Myoblast fusion requires several cell adhesion and transmembrane proteins, which accumulate at the contact sites between two myogenic cells. To further understand the role of Cap1, knockout and Cap1 overexpressing cells were incubated in the differentiation medium and relative mRNA levels of the profusion molecules integrin ß1D, caveolin, myomaker and myomixer were quantified by reverse transcription qRT-PCR. Interestingly, all profusion mRNA were significantly reduced in the Cap1 KO cells compared to the Cas9 control (Fig. 4B,C). In the case of Cap1 overexpressing cells, only the levels of integrin ß1D and myomixer decreased significantly (Fig. 4B,C). This indicates that the expression of profusion molecules is more severely affected by Cap1 knockout than by its overexpression, although the final result of myotube formation is impaired in both cases. Together, the results suggest that CAP1 perturbations impair differentiation by distinct mechanisms: whereas depletion hampers fusion, continuous overexpression of CAP1 affects myoblast elongation and alignment.

### Regulation of Cap1 expression by known myomiRs during differentiation

Since Cap1 is downregulated both at the mRNA and the protein level, we looked for the mechanism that might regulate the expression of Cap1 during differentiation. It is well established that the differentiation process of skeletal muscle involves miRNA called myomiRs which are upregulated manifold during differentiation (*27–30*). We thus revisited our own published dataset (GSE136956) (*27*) for differentiation-induced miRNA which might have potential binding sites in the 3’-UTR of Cap1. Based on predicted binding sites in the Cap1 mRNA we choose miR-1, miR-133 and miR-206, whereas miR-378 and miR-486 served as controls (no predicted binding site on the 3’-UTR of Cap1 mRNA) (Fig. 5A and B and Fig. S6).

**Fig. 5:**
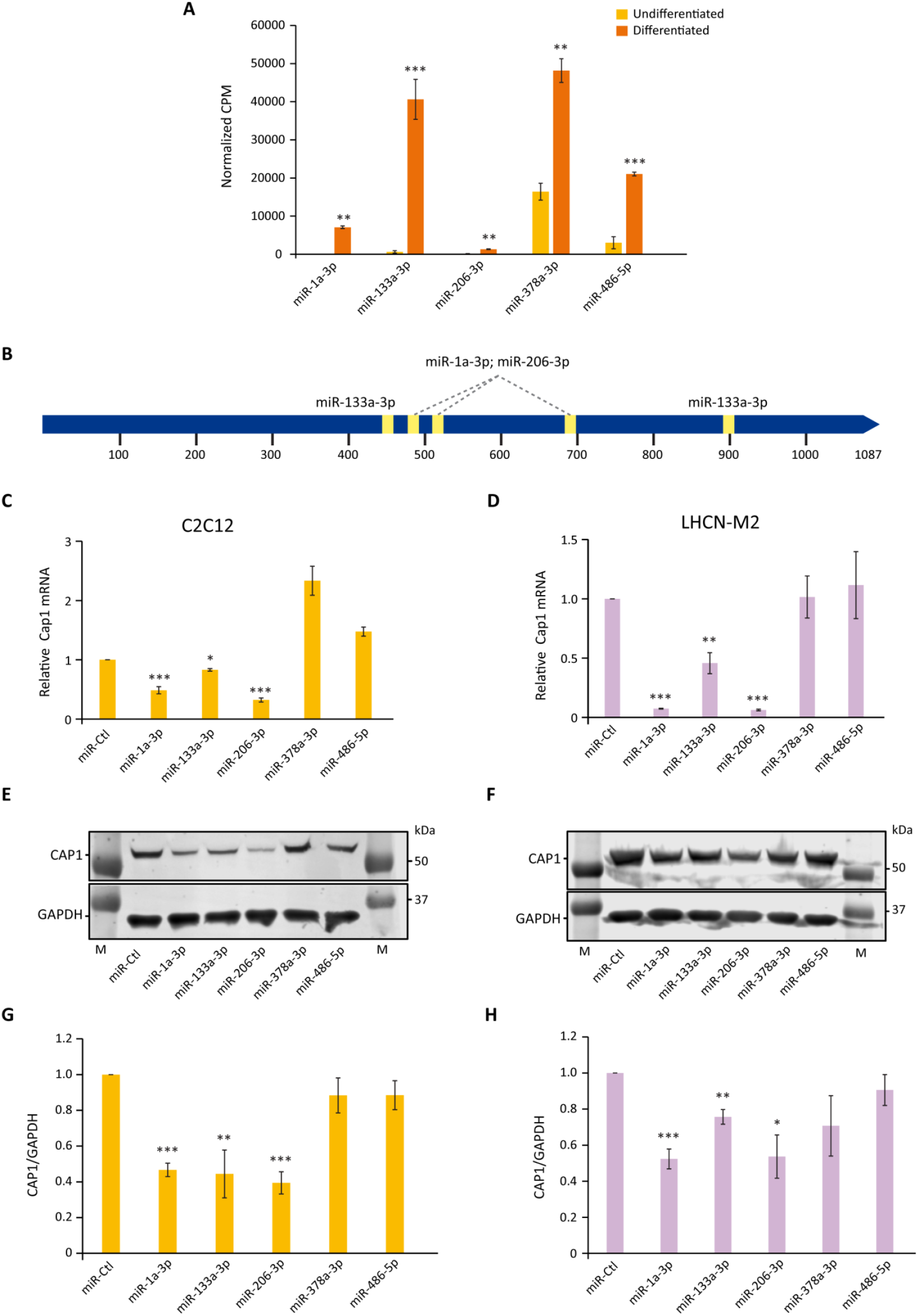
miRNA (miR-1, miR-133 and miR-206) regulate the expression of Cap1 in murine and human myoblast. (**A**) The abundance of the indicated miRNAs in total lysates of undifferentiated and differentiated C2C12, determined by RNA-Seq. CPM; counts per million. (**B**) Schematic of the 3′-UTR of murine Cap1 with the STOP-codon at position 1 and the polyadenylation signal at 1020 and 1058 bp. Predicted binding sites for miR-1, miR-133 and miR-206 are indicated by yellow boxes. (**C** and **D**) Cap1 mRNA expression in undifferentiated C2C12 (**C**) and LHCN-M2 (**D**) cells transfected with the indicated miRNA mimic for 72 h. (**E** and **F**) Representative immunoblots of cells transfected with the indicated miRNA. (**G** and **H**) Quantification of the CAP1 protein from three independent experiments. Error bars, SEM (n = 3); *p< 0.05, **p< 0.01, ***p< 0.001 (Student’s t-test).

Upon transfection of the corresponding miRNA mimics, Cap1 expression is downregulated by miR-1, miR-133 and miR-206 both at mRNA and protein level in C2C12 and LHCN-M2 cells (Fig. 5 C-H). In contrast, miRNAs which are shown to be upregulated during differentiation but lack predicted binding sites in the 3’ UTR did not downregulate the expression of the Cap1. The result suggests a posttranscriptional regulation of Cap1 expression by miRNAs, which is not secondary to the differentiation process per se.

### Cap1 reduction and myogenic differentiation requires post-transcriptional regulation via its 3’-UTR

To investigate whether Cap1 expression is directly regulated at the mRNA level, the 3’-UTR of the endogenous Cap1 locus was deleted by CRISPR–Cas9, using sgRNAs flanking the putative miRNA binding sites. Pools of infected and selected C2C12 cells were validated by PCR and sequencing of several cloned genomic loci, showing that the targeted region of the 3’-UTR between the STOP and the polyadenylation signal was successfully deleted (Fig. S7).

These ΔUTR C2C12 cells showed no reduction of Cap1 protein during differentiation, compared to the Cas9 control cells, which exhibited the significantly decreasing Cap1 expression observed before (Fig. 6A and B). In addition, myotube formation was impaired in cells with deleted Cap1 3’-UTR. The MYH marker of myogenic differentiation was reduced in the 3’-UTR-deleted cells, as shown for four and six days post differentiation (Fig. 6A and C). Morphologically, the ΔUTR C2C12 cells showed less mature, less multinucleated myotubes, compared to the Cas9 control cells (Fig. 6D). This appears to be reminiscent of the Cap1 overexpressing cells (Fig. 3 and Fig. S5). Together, these results indicate that posttranscriptional downregulation of Cap1 via its 3’-UTR is critical for myogenic differentiation, and suggest that a timely downregulation of Cap1 is important for controlling F-actin mediated changes of cell shape during maturation.

**Fig. 6:**
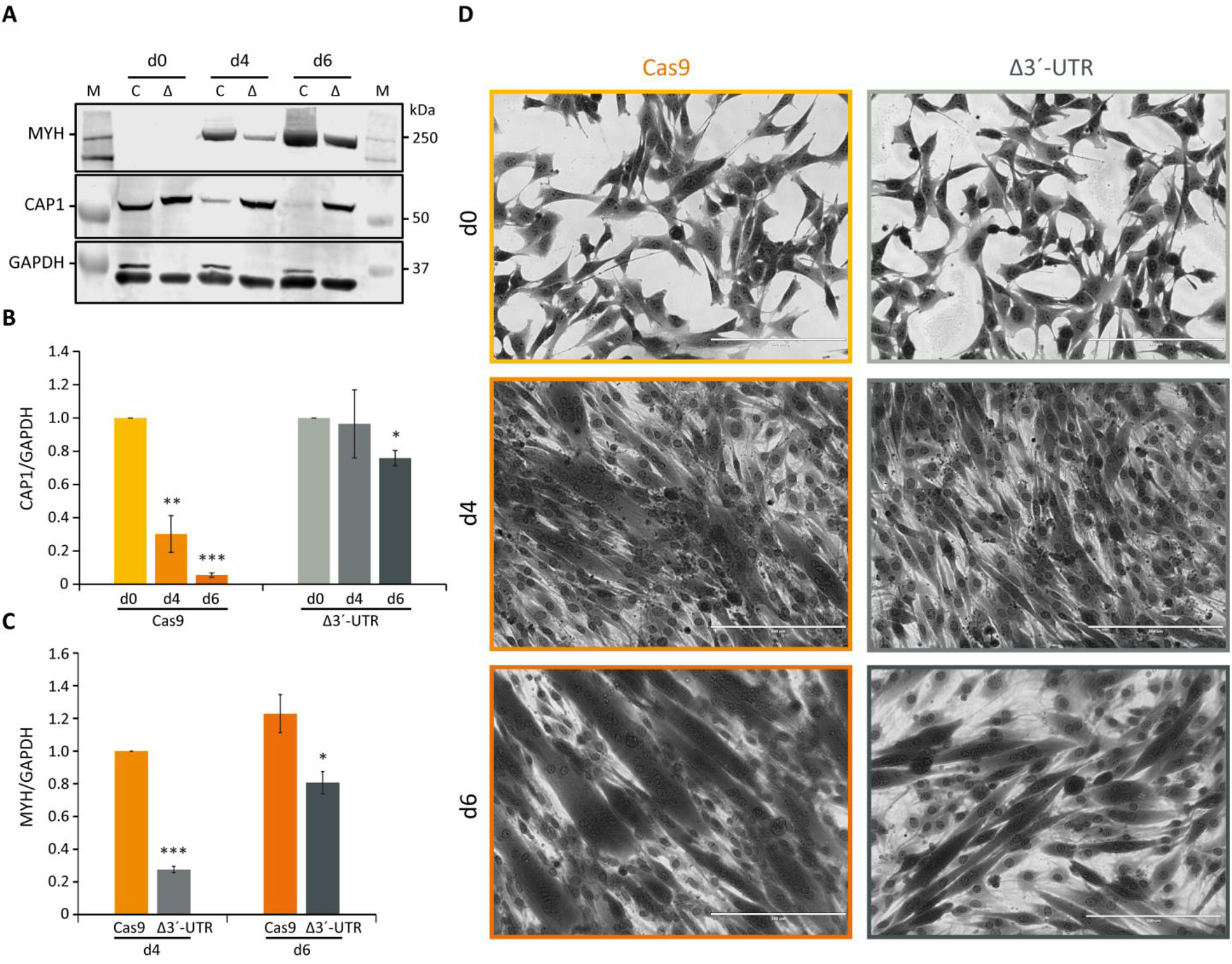
Requirement of the 3’-UTR for Cap1 regulation during myogenesis. (**A**) The 3’-UTR of Cap1 was deleted in C2C12 cells using CRISPR/Cas9. Δ3’-UTR (Δ) and Cas9-only (C) control cells were differentiated for the indicated times. Representative immunoblots for MYH and CAP1 are shown. (**B**) Quantification of CAP1 protein at day 0, day 4 and day 6 of differentiation normalized to day 0. (**C**) Quantification of MYH, normalized to Cas9 control cells at day 4. (**D**) Bright-field images (20x) of C2C12 cells stained with crystal violet after differentiation for 4 and 6 days (d4 and d6), in comparison to undifferentiated control cells (d0). Thick and multinucleated myotubes are reduced in the Δ3’-UTR cells (right panel) at day 6 of differentiation, compared to the Cas9 control (left panel). Error bars, SEM (n = 3); *p< 0.05, **p< 0.01, ***p< 0.001 (Student’s t-test). Scale bar, 200 μm.

**Fig. 7:**
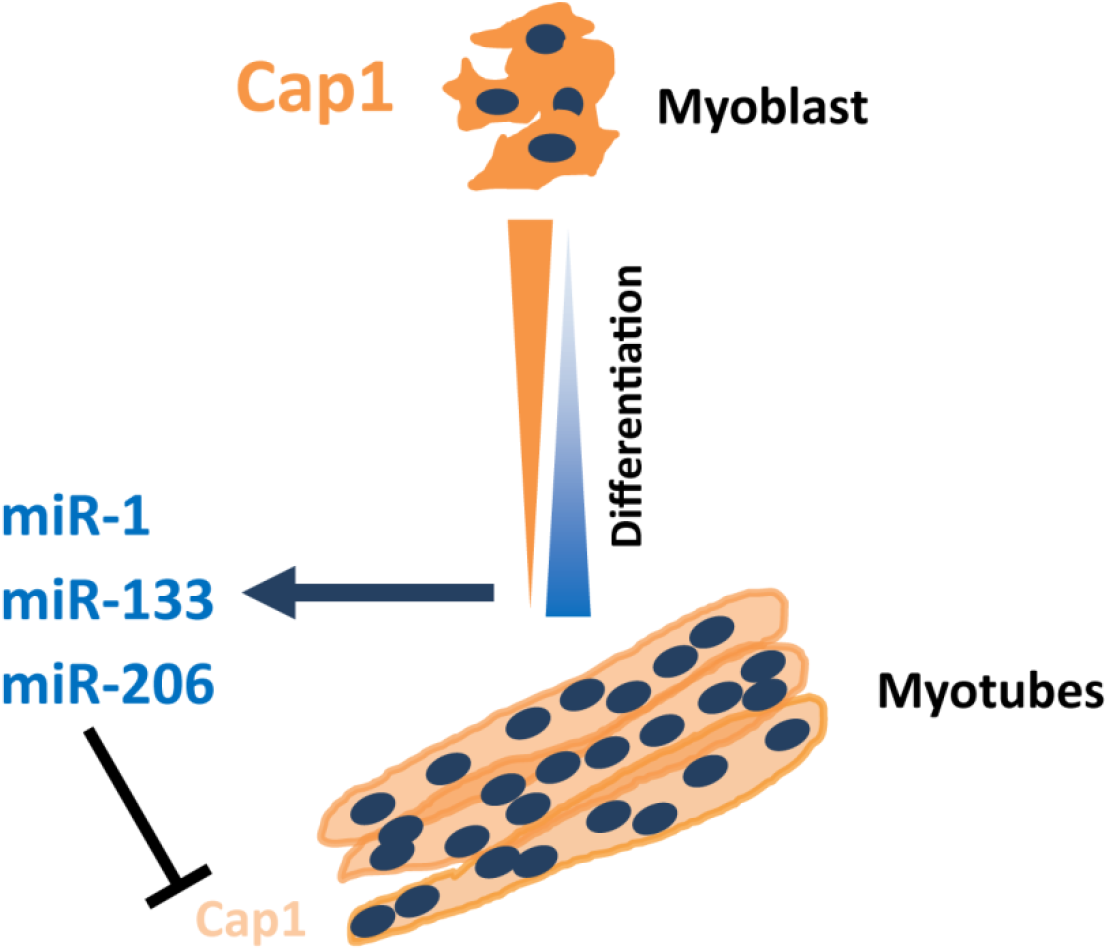
Model depicting the regulatory circuitry of myogenic C2C12 differentiation via post-transcriptional Cap1 regualtion. The timely and necessary downregulation of the Cap1 during myogenesis is induced by myogenic miRNAs miR-1a-3p, miR-133a-3p and miR-206-3p, which bind the Cap1 3′-UTR. miR-378a-3p and miR-486-5p, which do not have predicted binding sites on Cap1 3′-UTR do not regulate the expression of the Cap1, although, their expression also increases manifold during myogenesis. The decreased levels of Cap1 increases the F-actin levels which enables myoblasts for elongation, migration and fusion necessary for the myotube formation.

## Discussion

In the present study, we show a critical role of both Cap1 expression and it’s subsequent downregulation during myogenic differentiation. Our report implicates the miRNA-mediated downregulation of Cap1 in the myoblast-to-myotube differentiation. Myogenesis-induced upregulation of the myomiRs miR-1, miR-133 and miR-206 inhibit Cap1 expression, which in turn reduces the CAP1 protein amount. This enables the reorganization of F-actin that is required for changes in myoblast morphology and eventually the fusion process.

Cap1 is expressed in many tissues, but its physiological function remains largely unclear. Cap1 knockout mice do not survive post embryonic day 16.5 (*20*). A recent brain-specific Cap1 knockout shows a role in neuronal actin dynamics and growth cone morphology together with Cofilin1 (*21*). Cap1 in relation to skeletal muscle development or differentiation was not yet studied, probably due to almost undetectable levels of CAP1 in the adult skeletal muscle (*22, 23*). However, skeletal muscle Cap1 expression is detectable at the embryonic state, which reduces dramatically at birth and almost completely disappears in the adult. Similarly, Cap1 expression was found in microarray analysis of the skeletal muscle injury model, where Cap1 expression is upregulated immediately post-injury and disappears as the injury heals (*26*) (Fig. S2). Together with our data presented here, supports a role for Cap1 in muscle development and/or differentiation.

Both murine C2C12 and human LHCN-M2 are immortalized myoblast cells that partially recapitulate the myogenic differentiation *in vitro*. Both cells lines express Cap1, followed by a similar decrease upon induction of differentiation (Fig. 1), in agreement with microarray expression analysis previously performed by others (*25*) (Fig. S3). Thus, Cap1 might be expressed in activated skeletal muscle stem cell populations and myoblasts, while its expression decreases with maturation. In contrast, Cap2 is abundant in adult muscle and myotubes, and it is required for heart function and postnatal skeletal muscle development in mice and in human (*16, 17, 19, 31, 32*). Due to its abundant expression, most investigations of CAP proteins in muscle have thus focused on Cap2, but no roles in the early steps of myogenic differentiation have yet been characterized.

To date, a variety of experimental approaches have provided a basic framework of the processes that are required for myoblast fusion. On the cellular level, mammalian myoblasts seem to have an initial fibroblast-like morphology, which subsequently develops into an elongated, spindle-like shape capable of migrating towards other differentiating myoblasts (*9, 33*). Following contact, and membrane alignment, adhesion occurs between neighboring cells prior to membrane breakdown and fusion (*33–35*). During these early steps of differentiation, actin cytoskeleton rearrangements play a critical role. In a conditional Rac1 knockout mouse model, the recruitment of Arp2/3 and F-actin to the cell contact site is reduced, resulting in impaired myoblast migration and fusion (*36*). Experiments using C2C12 cells demonstrated that WAVE-dependent actin remodeling is required during myoblast fusion (*9*). In rat L6 myoblasts, the formation of a non-uniform cortical actin wall has been proposed, which provides structural and mechanical support to facilitate membrane alignment and fusion pore formation (*8*).

Concerning actin regulation, studies from the past decade have put CAP proteins at the center stage of the actin treadmilling regulation (*14, 15, 37, 38*). Towards this end, we could show that Cap1 reduction impaired the F-actin wall formation between C2C12 cells upon induction of the differentiation process (Fig. 4A). Rather, Cap1 knockout resulted in dramatically increased F-actin accumulations at uncoordinated places. On the contrary, overexpression of the Cap1 lead to a delayed response and cells still had a fibroblastoid shape, like the undifferentiated cell population (Fig. 4A). Together these data suggest that a timely upregulation in the F-actin content and its correctly localized arrangement upon induction of the differentiation plays a crucial role in the early steps of myoblast fusion.

After induction of differentiation, myoblasts exhibit an increased F/G-actin ratio, while the F-actin content of the cells decreases again at later stages and the ratio comes back to normal (*8*). This early increase in F-actin could be explained by the timely downregulation of CAP1, since the experimental reduction of Cap1 results in increased F-actin (Fig. 2) (*22, 39*). Further, an increased expression of Cap2 is observed during myogenic differentiation or regeneration, which potentially explains the normalization of the F/G-actin ratio towards completion of the process (Fig. S1–S3) (*25, 26*). Indeed, Cap2 upregulation during myogenesis is not essential for myoblast fusion or myogenic differentiation at the early stage, as evident from normal muscle development in Cap2 knockout mice, but for the late α-actin exchange during skeletal myofibril differentiation (*19*). Such a delay in the α-actin exchange has also been reported for mutant mice lacking Cofilin2, suggesting that CAP2 and Cofilin2 cooperate in myofibril differentiation (*40*). We speculate that Cap1 mediates actin regulation in the stem cell/myoblast compartment of the skeletal muscle and its downregulation increases appropriate F-actin structures needed for myoblast fusion, while the subsequent increase in Cap2 is needed to cut back the F-actin level and to play other important roles in myotube maturation.

Both overexpression as well as partial genetic knockout of Cap1 prevented proper differentiation of the C2C12 cells, as evident by a decrease in the signal for MYH in the western blot analysis (Fig. 3B, C). Interestingly, perturbation of Cap1 expression most prominently decreased myoblast fusion, as evident by the thickness and the number of nuclei per myotube (Fig. 3A and D). These results suggest that changes in the actin cytoskeleton by perturbing Cap1 expression primarily affect the fusion process. Although differentiation is similarly impaired, the excessive F-actin might be causative in the Cap1 knockout, while the lack of elongation may underlie the defects in Cap1 overexpressing cells (Fig. 4A). Such distinct reasons for a common outcome are similarly described for other actin modulator: Loss of Diaphanous results in less F-actin formation and reduced myoblast fusion, while its gain-of-function mutant massively increased F-actin foci, but ultimately also prevented myoblast fusion (*41*).

Our results show that Cap1 can regulate the expression of the myoblast profusion molecules ß1D-integrin, caveolin, myomaker and myomixer (Fig. 4B, C). This regulation must be indirect, as both knockout and overexpression of Cap1 resulted in their repression. Of note, the downregulation of these profusion molecules is more pronounced in Cap1 knockout cells as compared to overexpression. Thus there might be subtle differences in the mechanism which need further verification.

We identified miRNA-mediated mRNA degradation as a mechanism by which Cap1 is regulated. Both mRNA and protein levels were reduced during differentiation. It is well established that miRNAs play an important role during the myogenic differentiation process, and we and others have previously validated the upregulation of myomiRs whose expression increase manifold (*27–30*). We searched for the candidate miRNAs with potential binding sites in the 3’-UTR of Cap1, using our own published set of myomiRs in C2C12 cells (GSE136956) (*27*). Transient transfection of mir-1, mir-133 and miR-206 downregulate the expression of Cap1 both at mRNA and protein level, in contrast to the myomiRs mir-378 and miR-486 lacking an obvious 3’-UTR binding site (Fig. 5). Further, 3’-UTR sequence alignment from mouse, human, chicken, dog and rat show a conserved binding sites for the seed sequence of the tested miRNAs, indicating the conserved nature of the mechanism of Cap1 regulation (Fig. S6). Endogenous 3’-UTR deletion inhibited the downregulation of Cap1 and consequently impaired C2C12 differentiation, similar to the effects observed upon ectopic Cap1 expression (Fig. 6). At this point, however, other potential mechanisms of Cap1 regulation cannot be ruled out.

Taken together, Cap1 expression occurs in murine and human myoblasts and has an important role during myogenic differentiation. *In vivo*, muscle injury may activate quiescent stem cells such as satellite cells to form proliferating myoblasts, which profoundly express CAP1 protein. Moreover, timely downregulation of Cap1 by miRNA-mediated posttranscriptional control is required for upregulation of F-actin and further for the fusion process. Such a model is consistent with previous findings that Cap1 is expressed at the embryonic stage and upon injury, while its expression subsides upon healing and in the adult. We speculate that at later stages of the differentiation, Cap2 and probably other actin modulators compensate for Cap1 loss in myotubes. The mechanistic details and functional differences of these temporal changes, however, remain to be investigated.

## Materials and Methods

### Plasmids and reagents

Primers for qRT-PCR and sgRNA are listed in supplementary table 1. The cDNA clone for Cap1 was purchased from Origene (CAT#: MR207594) and subsequently cloned into lentiviral vector pLVX-puro (Clontech, Mountain View, USA). For overexpression of the candidate miRNAs, mirVana mimics (Thermo Fisher) were used together with RNAi-max (Thermo Fisher) following the manufacturer’s instructions.

### Cell Culture

C2C12 (DSMZ – German Collection of Microorganisms and Cell Cultures) was cultured subconfluently in Dulbecco’s modified Eagle’s medium (DMEM) supplemented with 10% fetal bovine serum (FBS), 2 mM L-glutamine, 1 mM sodium pyruvate, and antibiotic-antimycotic (Thermo Fisher) at 37° C and 5% CO_2_. Differentiation in C2C12 was induced by changing the medium to DMEM containing 2% horse serum (HS), 2 mM L-glutamine, 1 mM sodium pyruvate as well as antibiotic-antimycotic. Human immortalized LHCN-M2 myoblasts (Evercyte, Vienna, Austria; Cat. no. CkHT-040-231-2) were cultured in MyoUp medium (Evercyte, MHT-040). Following seeding, differentiation in the LHCN-M2 cells was induced by changing the medium (DMEM/M199 4:1, HEPES 20 mM, Zinc Sulfate 0.03 μg/ml, Vitamin B12 1.4 μg/ml, insulin 10 μg/ml and apotransferrin 100 μg/ml) and replacing it every second day.

For the generation of the Cap1 knockout (KO) C2C12 cells, two guide RNAs were used (Suppl. Table S1). The guide RNAs were designed by using the tool CHOPCHOP (*42*) (https://chopchop.cbu.uib.no/) and were chosen based on the fact they did not show any off-target predictions. Pools of knockout cells were validated for the specificity of the Cap1 deletion based on the downregulation in the Cap1. To generate the C2C12 cells carrying an endogenous deletion of the Cap1 3’-UTR, two guide RNAs (Suppl. Table S1) were designed to delete the majority of the 3 ‘-UTR from the Cap1 gene locus without affecting the coding region or the polyadenylation signal. sgRNA sequences were cloned into lentiCRISPR-V2 vector (Addgene; Plasmid # 52961, a gift from Feng Zhang) (*43*). C2C12 cells were lentivirally infected and selected with puromycin for one week. CRISPR and control pools of C2C12 cells were characterized for genetic deletion by PCR and sequence analysis. Generation of monoclonal cell lines is not possible because myoblasts have to be cultured subconfluently to maintain their undifferentiated state and fusion potential.

### Reverse transcription-quantitative PCR (qRT-PCR)

RNA was extracted using NucleoSpin RNA Mini kit for RNA purification (Macherey-Nagel Inc.) and cDNA was synthesized using Verso cDNA Synthesis Kit (Thermo Fisher Scientific), following the manufacturer’s instructions. Real-time PCR amplification and analysis were performed using a LightCycler 480 with Dynamo ColorFlash SYBR Green (Roche) and the primers are listed in Supplementary Table S1. For normalization, two different housekeeping mRNAs were used as controls. Calculations were done using the ΔΔ cycle threshold (Ct) method (*44*). For statistical analysis, the Student’s t-test was applied.

### Western blot analysis

Western blot analysis was performed following standard protocols. Primary antibodies against CAP1 (dilution: 1:500, Santa Cruz), Tubulin (1:5000, Sigma), Flag (1:1000, Sigma), Myosin Heavy chain 1/2/4/6 (1:1000, Santa Cruz), GAPDH (1:3000, Cell Signaling) were incubated overnight at 4°C. Fluorophore-conjugated secondary antibodies IRDye 700 or IRDye 800 (1:15000, LICOR Biosciences) were incubated for 1 h at room temperature. Imaging and quantifications were done using the Odyssey Image Scanner System with the software Image Studio V 3.1.4 (LI-COR Biosciences, Cambridge, UK), as described before (*45*).

### F-actin and crystal violet staining (cell size measurement)

For F-actin staining, cells were seeded in 12 well plates for 24 h, followed by differentiation as indicated. Cells were fixed with 3.7% PFA for 15 min. and permeabilized with 0.1% Triton X-100 for 10 min. After washing the cells were incubated with phalloidin (Alexa Fluor ™ 488 phalloidin, Thermo Scientific) in the blocking buffer following the manufacturer’s instruction. For crystal violet staining and cell size measurements, cells were seeded in a similar fashion at a sub-confluent level in 12 well, plate, and post 24 h they were incubated with 0.5% crystal violet in methanol for 20 min. The cell size measurement was done by ImageJ software.

### Immunofluorescence microscopy for myosin heavy chain

For immunofluorescence microscopy cells were fixed with 3.7% formaldehyde, permeabilized with 0.2% Triton X-100 and blocked with 10% horse serum (Sigma Aldrich, München, Germany), 1% BSA (Carl Roth), 0.05% Triton X-100 (Carl Roth) in PBS. The following antibodies were used: Myosin Heavy chain 1/2/4/6 (1:500, Santa Cruz) and secondary antibodies conjugated with Alexa488 (1:500, Thermo Fisher Scientific). DNA was counterstained with DAPI (Sigma Aldrich). Samples were covered with Immu-Mount (Thermo Fisher Scientific) and imaged with an Axio Observer 7 (Zeiss, Jena, Germany) equipped with a monochrome Axiocam.

### Transient transfection of the miRNA mimics and Western blot analysis

Both the C2C12 and LHCN-M2 cells were transiently transfected with miRNA mimics (mirVana mimics, Thermo Fisher). Briefly, the cells were seeded at sub-confluent density in the 6 well plate format and were transfected with 50 nmoles of miR-Ctl, miR-1a-3p, miR-133a-3p, miR-206-3p, miR-378-3p, or miR-486-5p mimics using 10 μl of RNAimax following the manufacturer’s instruction. After 72 h, cells were lysed and analyzed for the effect of the miRNA transfection on Cap1 mRNA and protein level by reverse transcription quantitate PCR and western blot analysis respectively, as described above.

## Acknowledgments

We thank Ines Block and Igor Kovecevic for critically reading of the manuscript.

## Author contributions

AKS conceived and designed the study. AKS and AW conducted the experiment and AKS analyzed the data. AR did the cloning of the over-expression constructs, helped with figure preparation and analyzed the result. GP analyzed the data and supervised the study. GP, AKS and AR wrote the manuscript. All the authors read and approved the final manuscript

## Data and materials availability

All data needed to evaluate the conclusions in the paper are present in the paper main text and/or the supplementary materials.

## Supplementary Materials

**Fig. S1:**
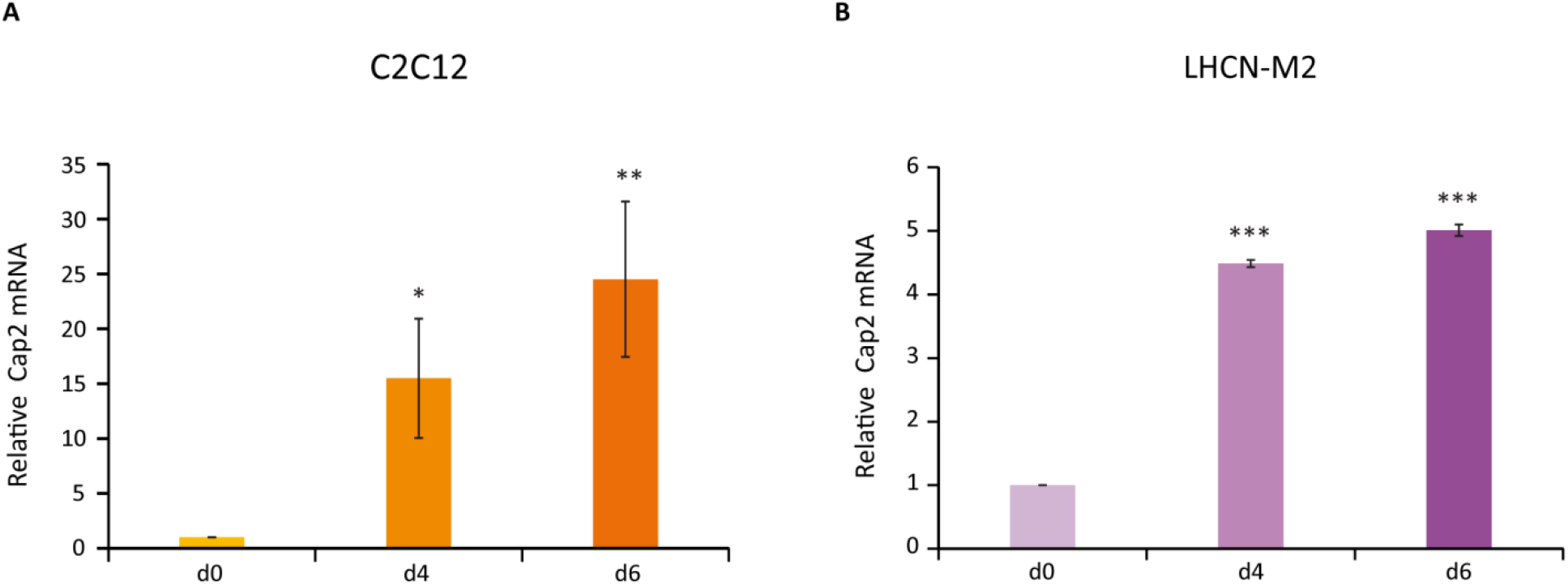
Cap2 mRNA increases during myogenic differentiation of the murine and human myoblast cell lines. (**A**) Expression of the Cap2 increases significantly in the C2C12 cells during differentiation day 4 and day 6 (d4 and d6) as compared to the undifferentiated control cells (d0). (**B**) A similar increase in the Cap2 mRNA level was observed in the case of human myoblast cell line LHCN-M2. Error bars, SEM (n = 3); *p< 0.05, **p< 0.01, ***p< 0.001 (Student’s t-test).

**Fig. S2:**
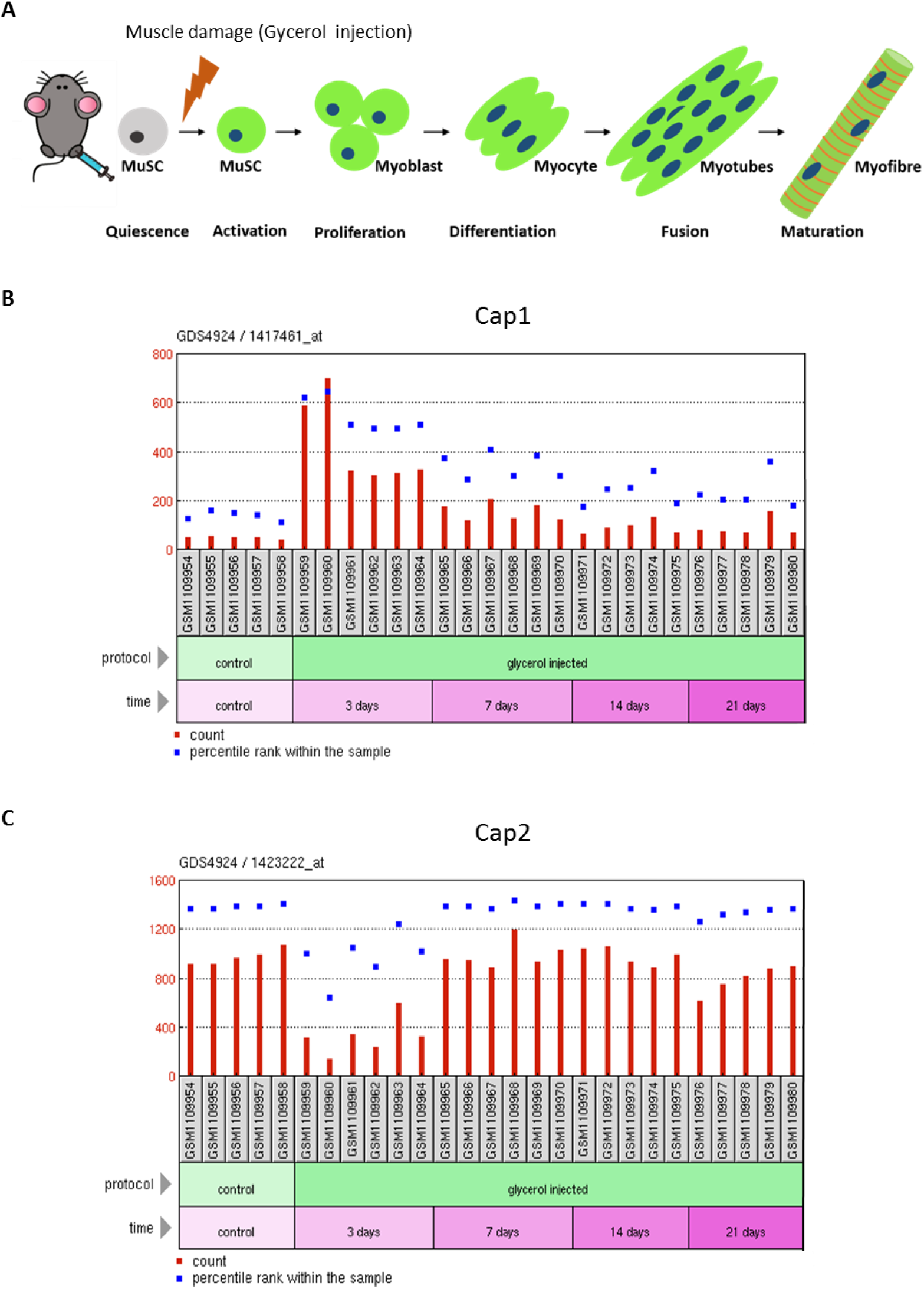
Re-analysis of GEO dataset GSE45577, showing induction of Cap1 expression after injury of skeletal muscle *in vivo*. (**A**) Schematics of the experimental setup for the mouse skeletal muscle injury model by (*1*). (**B**) Expression of Cap1 in the tibialis anterior muscle of 12 wk old C57BL/6J males before and after injection of glycerol, using the Affymetrix Mouse Genome 430 2.0 Array. Significant upregulation/re-expression of Cap1 occurs 3 days post-injury, followed by subsequent downregulation towards 21 days when regenerated mature myotubes show only diminished Cap1 expression (https://www.ncbi.nlm.nih.gov/geoprofiles/113963492). (**C**) Cap2 expression showing inverse regulation upon regeneration, with low-level expression in muscle 3 days after injury (https://www.ncbi.nlm.nih.gov/geoprofiles/113969228).

**Fig. S3:**
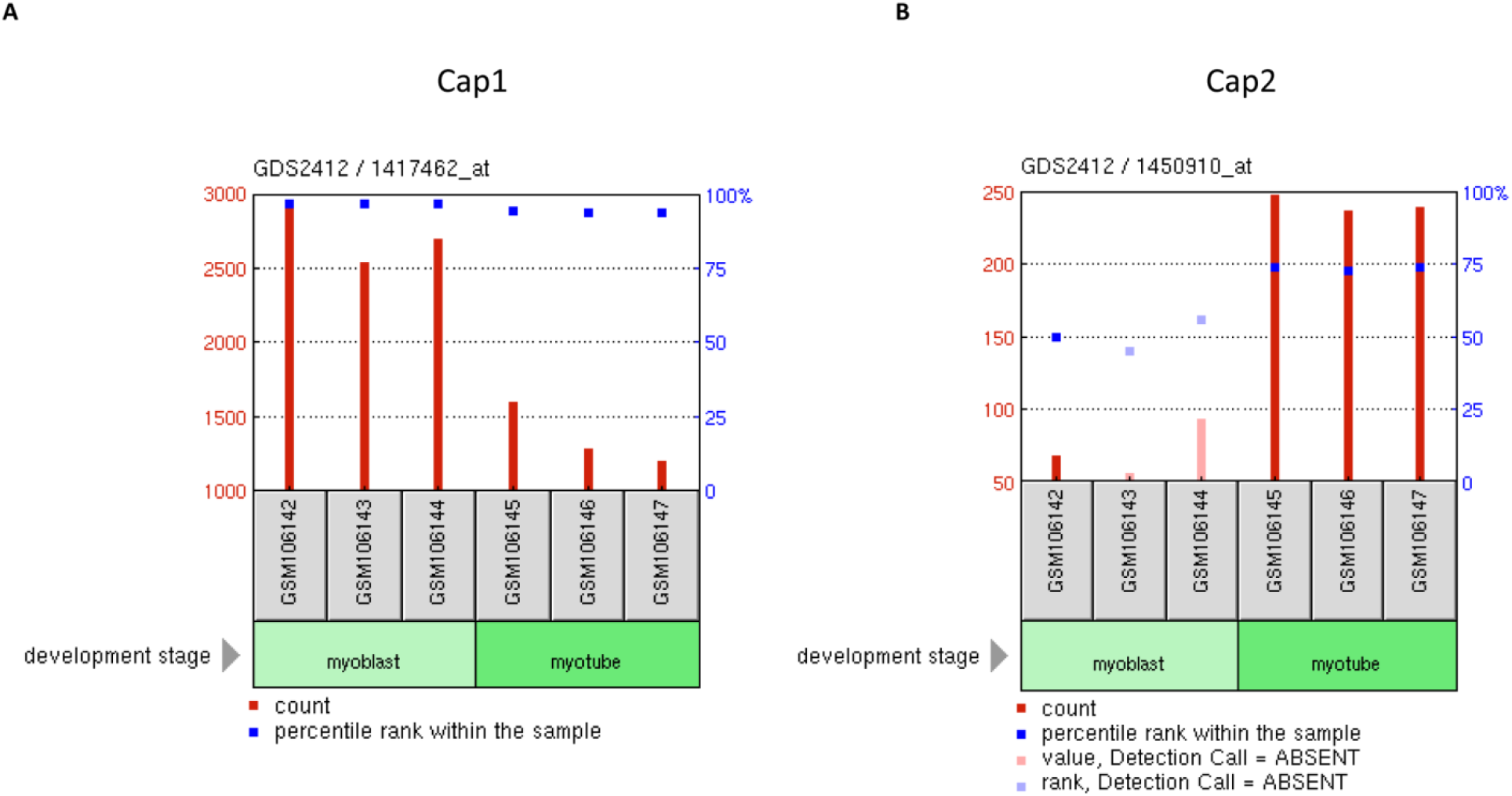
Re-analysis of GEO dataset GSE4694, showing diminished Cap1 expression upon myogenic differentiation of C2C12 cells. (**A**) Expression of Cap1 in the Affymetrix Mouse Genome 430 2.0 microarray by (*2*). Compared to the myoblast population of C2C12 cells, differentiated myotubes show diminished expression of Cap1 (https://www.ncbi.nlm.nih.gov/geoprofiles/31829693). (**B**) Inverse regulation of Cap2, being almost undetectable in undifferentiated C2C12 myoblasts (https://www.ncbi.nlm.nih.gov/geoprofiles/31863105).

**Fig. S4:**
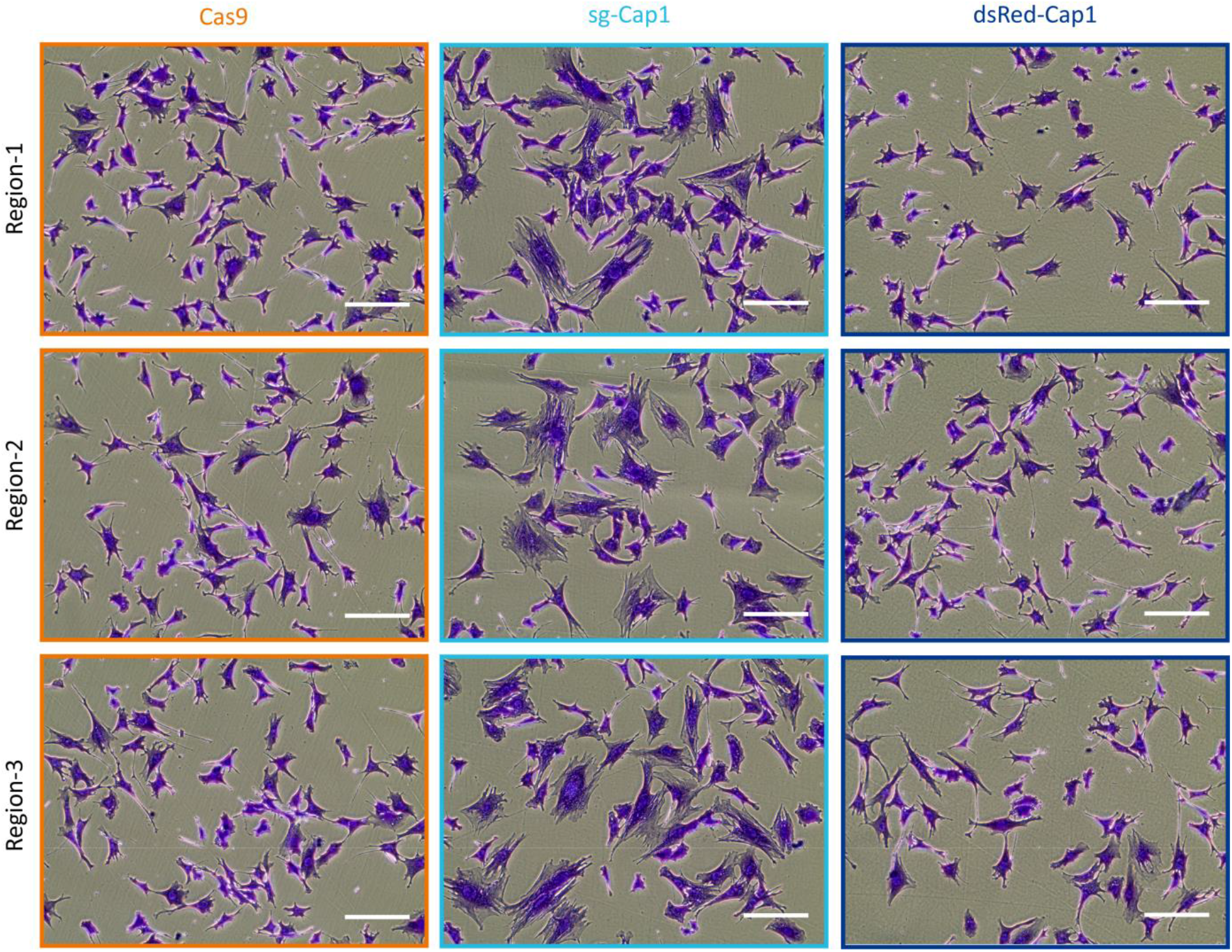
Knockout (KO) of Cap1 resulted in increased cell size. Crystal violet staining of the CRISPR-Cas9 mediated Cap1 KO (sg-Cap1; mixed population, middle panel), shows a much bigger cell size as compared to the Cas9 control cells (left panel). While overexpression (OE) of the tagged Cap1 (dsRed-Cap1) did not affect the cell size (right panel). Microscopic pictures from three different areas are presented in the figure. Scale bar, 200 μm.

**Fig. S5:**
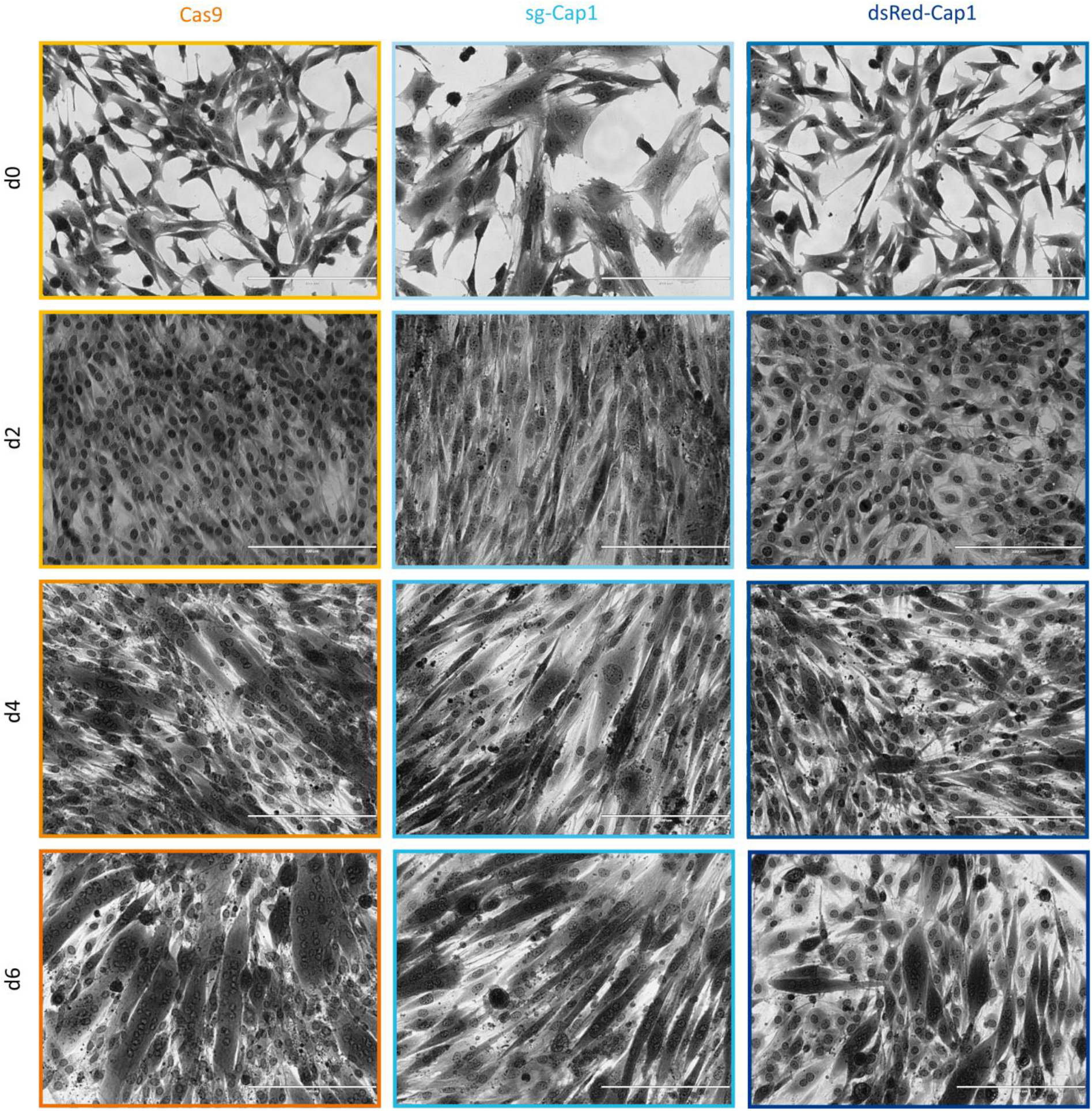
Perturbation in Cap1 expression affects the myogenic differentiation of C2C12 cells. Bright-field images (20x) of C2C12 cells upon differentiation for 2, 4 and 6 days (d2, d4, d6), in comparison to undifferentiated control cells (d0). On day2 (d2) of the differentiation, both the Cas9 controls as well as the KO cells (sg-Cap1) show a more elongated phenotype, while Cap1 OE cells (dsRed-Cap1), still showed more fibroblast (parental) phenotype. Day4 (d4), of the differentiation, shows the appearance of multinucleated myotubes in the case of the Cas9 cells, while in the case of the KO as well as OE cells they appear to have a single nucleus per cell. Cap1 over-expressing cells show now elongated phenotype on the day4 of the differentiation. On day6 (d6) of the differentiation Cas9 cells showed more mature myotubes with more numbers of nuclei per myotube as compared to both KO and OE cells line, which also showed the multinucleated myotubes but the number of nuclei per myotubes was significantly low. Scale bar, 200 μm.

**Fig. S6:**
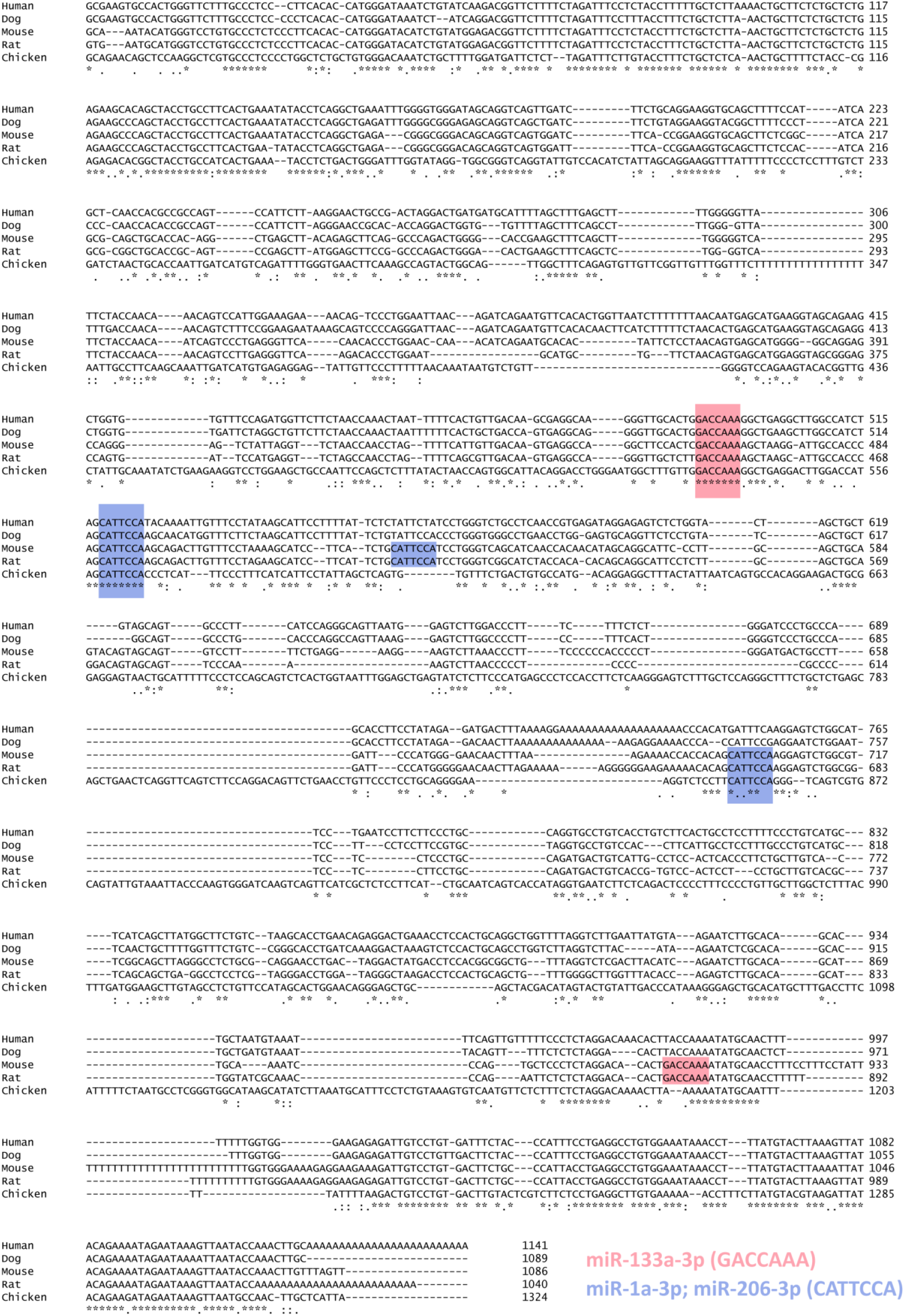
Multiple sequence alignment (*3*) of Cap1 3’ UTR from different organisms shows conserved seed sequences for the tested miRNA. Red boxes indicate the seed sequence for the binding site for miR-133a-3p, and blue boxes for the miR-1a-3p and miR-206-3p, which share the same seed sequences.

**Fig. S7:**
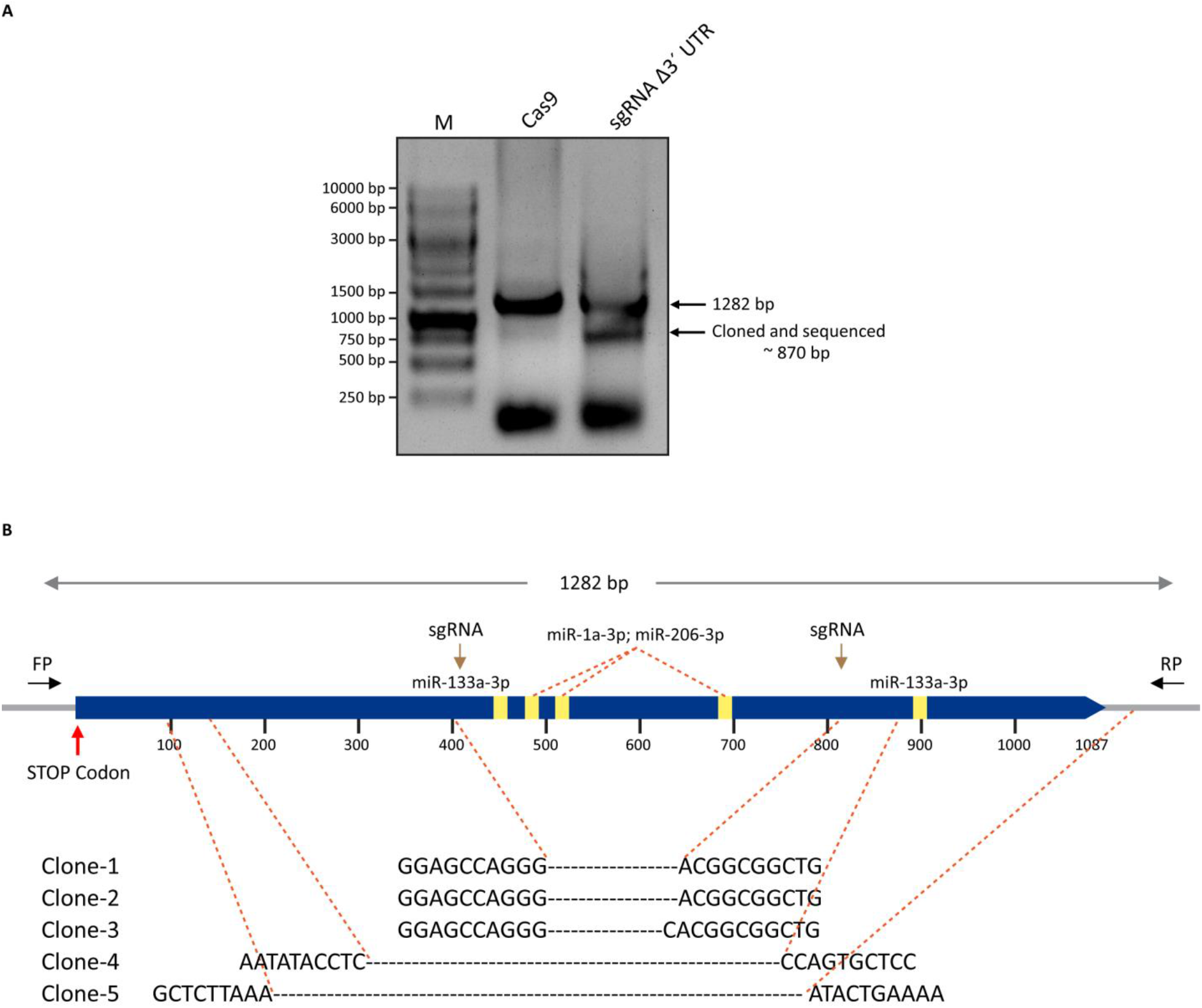
CRISPR-Cas9 mediated deletion of 3’-UTR of Cap1 endogenous locus. (**A**) DNA-Agarose gel from the PCR primers depicted in the schematic diagram. The appearance of the smaller fragment represents the cell population with the deleted part of the 3’-UTR at the endogenous locus (mixed population). The smaller PCR amplicon from the gel was cloned and sent for sequencing to confirm the deletion. (**B**) Schematic of the 3′-UTR of murine Cap1 with miRNA binding sites indicated by yellow boxes and sgRNA target sites indicated by an arrow and forward and reverse primer binding position corresponding to the genomic DNA.

**Table S1:**
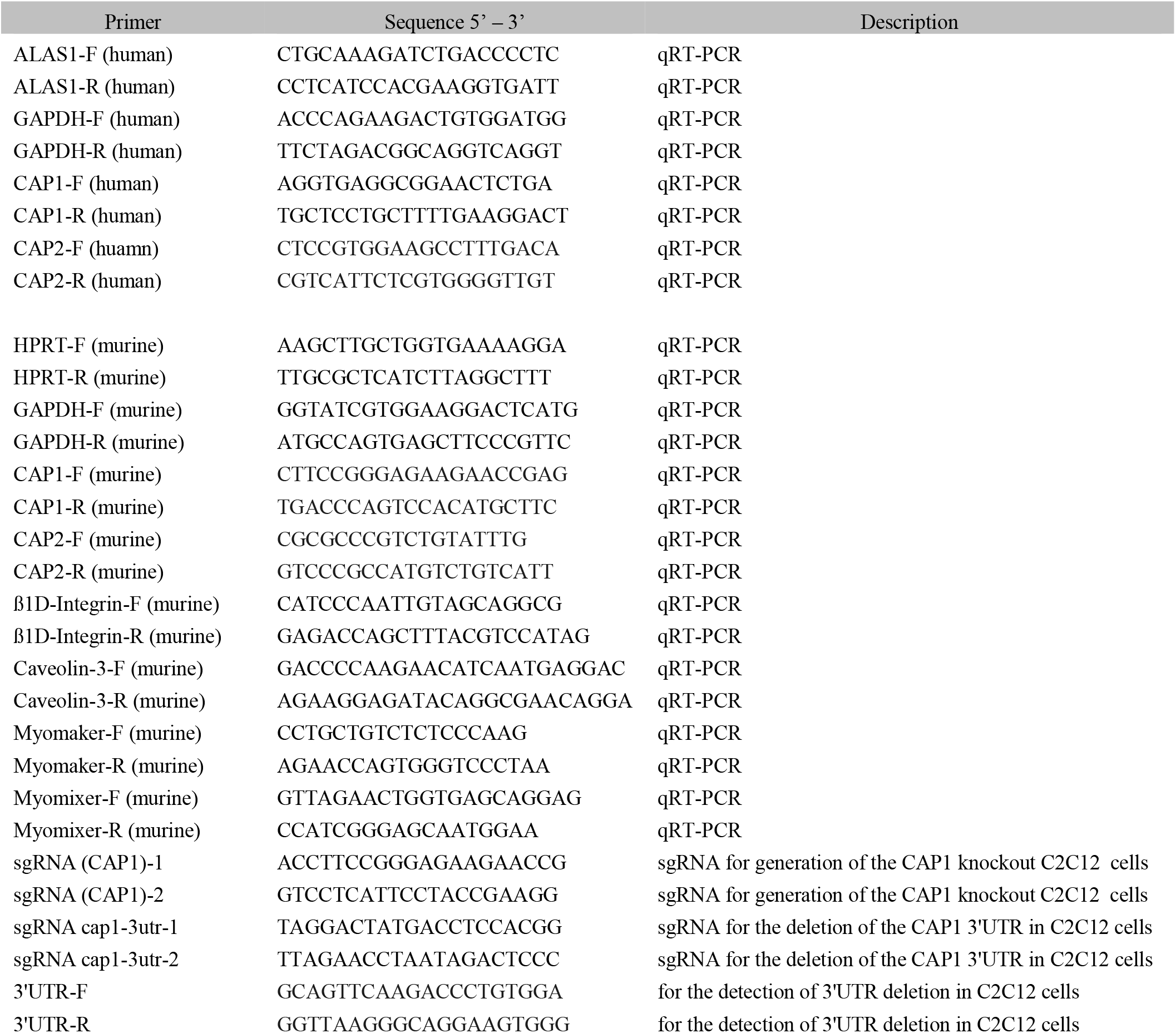
Primer list used for the generation and detection of CRISPR/Cas9 KO cells and quantitative real-time PCR.

